# A distinct phylogenetic cluster of Monkeypox genomes suggests an early and cryptic spread of the virus

**DOI:** 10.1101/2022.07.30.502168

**Authors:** Bani Jolly, Vinod Scaria

## Abstract

Since its first reports in humans in 1970, monkeypox has been predominantly restricted to countries in Africa where the disease is endemic. Early in 2022, a large number of cases of the disease were reported from Europe and other countries in patients with no history of travel to regions where monkeypox is endemic. Amid a rise in cases, the availability of genome sequences of monkeypox virus isolates in the public domain provides an opportunity to understand the transmission and evolution of the virus. Here, we describe a distinct phylogenetic cluster of monkeypox virus (lineage A.2) using genome sequences available on GISAID. Lineage A.2 currently encompasses 9 genome sequences from 6 viral isolates collected from 3 countries and is distinctly different from the predominant lineage B.1 which is linked to the large European outbreak.

---

Monkeypox, a rare viral infection, has been predominantly restricted to countries in Africa and was infrequently reported from regions where the disease is not endemic, until recently. Early in 2022, an outbreak of the disease has been reported and has been epidemiologically linked to potential superspreader events in Europe earlier in the year^1^. Over a short period, the disease has spread across 75 countries causing over 20,000 infections, as on 30 July 2022, with the World Health Organization subsequently declaring the outbreak as a public health emergency of international concern in July 2022^2^. Following the outbreak, the wide availability of monkeypox genome sequences in the public domain provided a unique opportunity to understand the genetic epidemiology as well as the evolution of the pathogen.

With heightened surveillance and molecular diagnosis across the globe, several genomes of the monkeypox virus are being deposited in public databases like GISAID^3^. As early epidemiologic studies linked the outbreak of monkeypox in 2022 to superspreader event(s) in Europe, it was therefore not surprising that a majority of the genomes available on GISAID cluster together. Further reports suggest that the early transmission in this outbreak was largely among gays, bisexuals and other men who have sex with men (MSM) with some exceptions^4,5^. However, the recently deposited genome sequences from the United States of America, Thailand and India suggest a distinctly different phylogenetic cluster of genomes, classified by Nextclade as lineage A.2, in contrast to the cluster encompassing the majority of genomes (N=547) that are classified as lineage B.1 (**Figure 1A**)^6–8^. The A.2 lineage comprises 9 genomes from 6 unique clinical isolates, with the earliest genome belonging to this lineage (collected in July 2021) being deposited from the state of Texas, USA. The two genome isolates from India cluster closely with a genome isolate from Florida (hMpxV/USA/FL-DHCPPCDC-001/2022) on the phylogenetic tree, while the isolates from Texas, Virginia and Thailand mapped to separate sub-clusters (**Figure 1B**).

**Figure 1.**
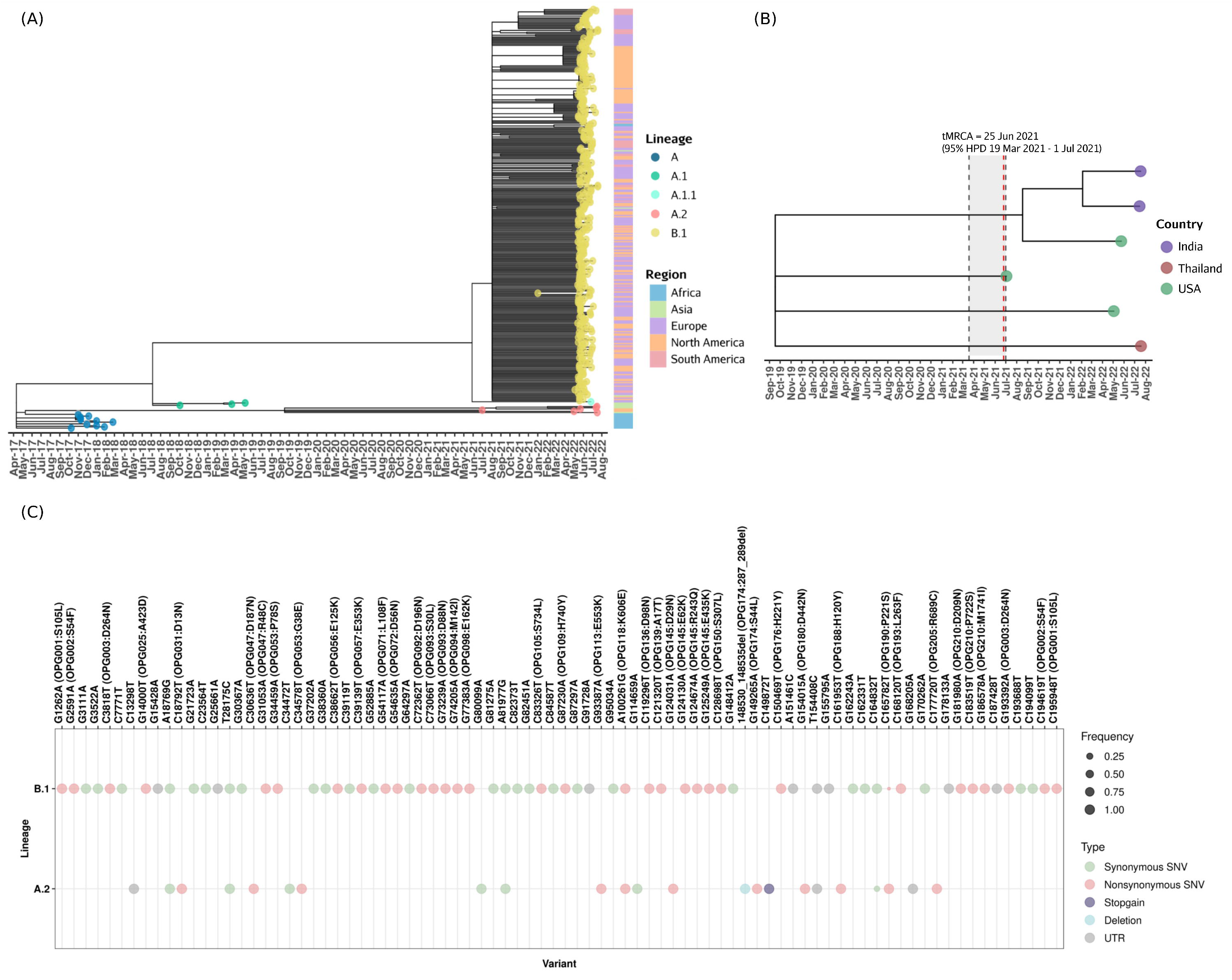
**(A)** Phylogenetic tree of genome isolates from GISAID belonging to the hMPXV-1 clade of monkeypox virus constructed using Nextstrain^6^. **(B)** Phylogenetic tree of the genomes belonging to A.2 monkeypox lineage. **(C)** Comparison of frequencies of variations present in >90% genomes of lineages A.2 (N=6) and B.1 (N=547).

The genomes belonging to the A.2 lineage have 16 distinct genetic variations which are not found in other lineages, of which 9 are nonsynonymous, 3 are synonymous and 1 is a stopgain variation, along with a deletion of 3-amino acid deletion in the gene OPG174 (**Figure 1C**). Variants found at a minimum frequency of 90% in the respective lineages A.2 and B.1 were compared and are summarised in **Figure 1C**.

Albeit the limited number of sequences available for the A.2 lineage, we attempted to compute the time to the most recent common ancestor (tMRCA) for A.2 and the nucleotide substitution rates for lineages A.2 and B.1 using BEAST v1.10.4^7^. The tMRCA was calculated following a coalescent growth rate model with a strict molecular clock and the HKY+Γsubstitution model. MCMC was run for 50 million steps and the initial 1% steps were discarded as burn-in. The tMRCA of the A.2 lineage was computed as 25 June 2021 (95% HPD 19 March 2021 to 1 July 2021). The A.2 lineage had a mean nucleotide substitution rate of 5.53×10^−5^ (95% HPD 3.39×10^−5^ to 7.46×10^−5^) substitutions per base/year, suggesting a modest rate of substitution compared to the substitution rate of 1.13×10^−4^ (95% HPD 9.33×10^−5^ to 1.33×10^−4^) substitutions per base/year for the larger B.1 lineage of genomes. Accelerated evolution of the B.1 lineage has been observed recently^8^.

Limited demographic information could be linked to the members of the A.2 cluster and has been primarily compiled from the metadata associated with the genome sequences and reports in the public domain. The two genomes from Kerala, India were isolated from men who had a travel history to the United Arab Emirates, while the genome from Thailand was isolated from a male traveller from Nigeria. The genome from Texas, United States of America, the earliest in the cluster, was also isolated from a male traveller from Nigeria suggesting a wider geographic area with ongoing transmission of the virus beyond regions in Central and Eastern Africa where the virus is endemic^9^.

Put together, the evidence suggests that this unique and distinct phylogenetic cluster of genomes, therefore, represents sustained and previously uncharacterized human-human transmission events spanning multiple countries. The tMRCA dating to mid-2021 and the earliest genome dating to July 2021 suggests that this sustained transmission event possibly preceded the outbreak in 2022 in Europe and has remained largely undetected. The distinct genomic signatures suggest that this transmission chain may not be linked to the large outbreak of monkeypox which occurred in 2022 and has been potentially uncovered due to heightened awareness, surveillance and the wider availability of diagnostics.

This report, therefore, re-affirms the unique and significant value of genomic surveillance of emerging pathogens in uncovering potential new insights and leads for epidemiological investigations. The distinctive finding in this report may have a significant impact on public health policies, surveillance as well as public-health communication.

## Acknowledgement

BJ acknowledges research fellowships from the Council of Scientific and Industrial Research (CSIR), India. The authors acknowledge Division of High Consequence Pathogens and Pathology (DHCPP, CDC) - the United States of America, Indian Council of Medical Research (ICMR), National Institute of Virology (NIV) - India, National Institute of Health - Thailand, Thai Red Cross Emerging Infectious Diseases Clinical Center, Faculty of Medicine (Chulalongkorn University) - Thailand, and other investigators for the data deposited in the public domain. A detailed list of acknowledgements is available at https://github.com/banijolly/Phylovis-MPX. The authors also acknowledge Dr. Sandhya Pulukool and Vishu Gupta for offering suggestions and insights for enriching the manuscript.

